# Differential impact of acute and chronic stress on CA1 spatial coding and gamma oscillations

**DOI:** 10.1101/2021.04.23.441067

**Authors:** Anupratap Tomar, Denis Polygalov, Thomas McHugh

## Abstract

Chronic and acute stress differentially affect behaviour, as well as the structural integrity of the hippocampus, a key brain region involved in cognition and memory. However, it remains unclear if and how the facilitatory effects of acute stress on hippocampal information coding are disrupted as the stress becomes chronic. To examine this, we compared the impact of acute and chronic stress on neural activity in the CA1 subregion of male mice subjected to a chronic immobilization stress paradigm. We observed that following first exposure to stress (acute stress), the spatial information encoded in the hippocampus sharpened and the neurons became increasingly tuned to the underlying theta oscillation in the local field potential (LFP). However, following repeated exposure to same stress (chronic stress), spatial tuning was poorer and the power of both the slow-gamma (30-50 Hz) and fast-gamma (55-90 Hz) oscillations, which correlate with excitatory inputs into the region, decreased. These results support the idea that acute and chronic stress differentially affect neural computations carried out by hippocampal circuits and suggest that acute stress may improve cognitive processing.

## Introduction

It is generally accepted that while mild or acute stress can be beneficial for cognition and learning, repeated exposure to stressors (chronic stress) disrupts these processes (Luksys and Sandi, 2011). This dichotomy in the impact of acute and chronic stress has also been observed in the hippocampus, a brain region crucial for acquisition and consolidation of declarative memory. At the cellular level, chronic, but not acute stress, causes dendritic shrinkage and debranching (Sousa et al., 2000; Watanabe et al., 1992) and decreases number of synaptic contacts (spines) on principal hippocampal pyramidal neurons (Magariños et al., 1997; Sandi et al., 2003). Further, earlier studies employing both *ex vivo* electrophysiology and *in vivo* tetrode recordings report that chronic stress also alters the functionality of hippocampal pyramidal cells. For example, chronic stress disrupts synaptic plasticity in hippocampus slices (Alfarez et al., 2003). Similarly, the spatial map, or internal representations of surroundings (O’Keefe and Nadel, 1978), evident in the location-specific increase in average firing rate of hippocampal pyramidal “place” cells (O’Keefe, 1976; O’Keefe and Dostrovsky, 1971) is altered in chronically stressed rodents (Kim et al., 2007; Passecker et al., 2011; Tomar et al., 2015). However, the interpretation of acute stress effects on hippocampal synaptic plasticity are more complex (Joëls and Krugers, 2007; MacDougall and Howland, 2013) and consequently, the impact of acute stress on the neural computations carried out by hippocampal circuits in the intact brain remains unclear.

In addition to the rate code (i.e. location-specific spiking), place cells also use temporal coding to signal spatial aspects of the animal’s location or behaviour (O’Keefe and Recce, 1993). Temporal coding involves place cells spiking at a specific phase of ongoing oscillations in the local field potential (LFP), such as theta (6-12 Hz) and gamma (30-90 Hz), during exploratory behaviour. These oscillations, as well as the more transient coupling of the theta-gamma oscillations themselves, are thought to provide temporal precision to the activity of hippocampal cell assemblies and to facilitate phenomena including synaptic plasticity and retrospective and prospective coding (Buzsáki, 2010; Fries, 2015; Harris et al., 2003; Lisman, 2005). Interestingly, temporal coding, as well as theta and gamma coupling, has been shown to be altered in neurodegenerative disorders (Booth et al., 2016; Goutagny et al., 2013; Mably et al., 2017), for which stress is a risk factor (Bisht et al., 2018). Thus, it is likely that both acute and chronic stress may impact these oscillatory patterns in unique ways.

To address these gaps in our knowledge we employed tetrode recordings in the dorsal CA1 of male mice. Recordings were made while mice explored a linear track before and after experiencing chronic immobilization stress (CIS) (Suvrathan et al., 2010), a protocol that has been previously shown to reduce hippocampal volume, spatial memory (Rahman et al., 2016) and context discrimination (Tomar et al., 2015). Specifically, we examined alterations in both rate and temporal coding of CA1 pyramidal cells, as well as changes in the hippocampal oscillatory activity, following acute and chronic stress.

## Results

The main aim of this study was to examine the differential impact of acute and chronic stress on CA1 spatial coding and hippocampal physiology. To this end, we employed a longitudinal design similar to that employed in previous studies that contrasted neural activity from same rodents before and after they received exposure to stress (Ghosh et al., 2013; Kim et al., 2007; Tomar et al., 2021). Specifically, we monitored CA1 place cell activity and theta (6-12 Hz) and gamma oscillations (30-90 Hz) during two track exploration sessions: one occurring before (PRE) and second after (POST) the stress exposure, on the first day (Acute) and the last day (Chronic) of a chronic immobilization stress (CIS) paradigm (Fig. 1A; see Methods), thus providing us with four conditions, i) PRE-Acute, ii) POST-Acute, iii) PRE-Chronic iv) POST-Chronic.

**Fig. 1.**
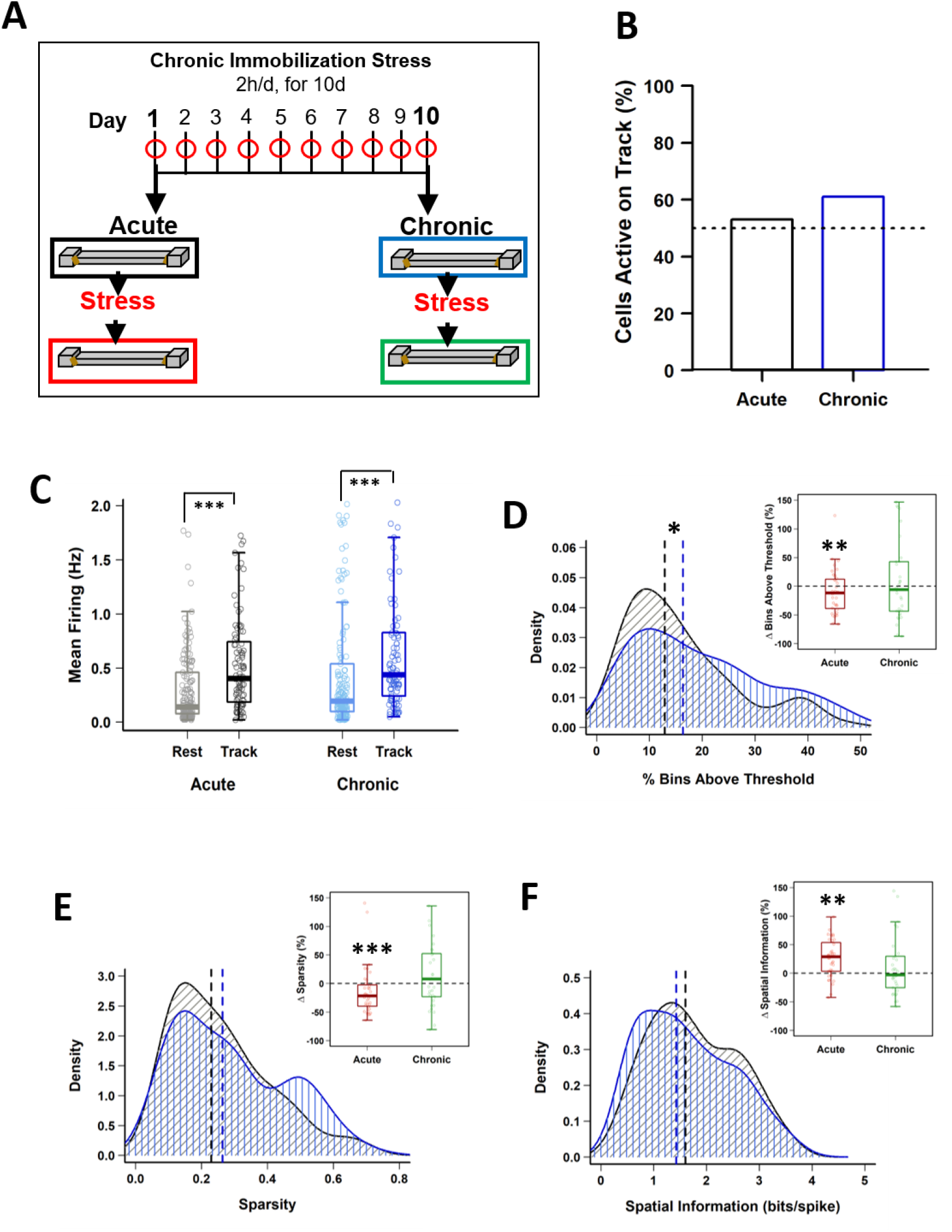
Impact of stress on CA1 place cell activity. (A) Schematic representation of the chronic immobilization stress (CIS) protocol and experimental design. (B) Percentage of pyramidal cells active during exploration (RUN) compared to quiet wakefulness/sleep (REST) period, before stress administration (Pre-stress), on day-1(black) and day-10 (blue), (day-1: 95/180 (53%) vs day-10: 101/166 (61%), p = 0.164, chi-square test). Dotted line represents 50%. (C) Pre-stress, mean firing rate between REST and RUN on day-1 and day-10 (LMMs: main effect of day, F_(1, 538)_ = 3.988, p = 0.046; main effect of session, F_(1, 538)_ = 64.02, p = 7.708 × 10^−15^; interaction, F_(1, 538)_ = 0.018, p = 0.892; post-hoc Tukey’s test, day-1: REST vs RUN, p<0.0001, day-10: REST vs RUN, p < 0.0001). (D) Pre-stress place field size density distribution differs between day-1 and day-10 (PRE-Acute vs PRE-Chronic, p=0.049, KS-test,). However, place cells active during RUN, before and after stress exposure, display a decrease in field size on day-1 (PRE-Acute, 13.89 ± 1.46 vs POST-Acute, 11.25 ± 0.96, V=280, p= 0.009, Wilcoxon signed-rank test) but not on day-10 (PRE-Chronic, 19.71 ± 2.39 vs POST-Chronic, 17.81 ± 1.99, V = 186, p = 0.316, Wilcoxon signed-rank test). (E) Pre-stress sparsity of place fields does not differ between day-1 and day-10 (PRE-Acute vs PRE-Chronic, p>0.05, KS-test). However, place cells active during RUN, before and after stress exposure, display a decrease in sparsity-index on day-1 (PRE-Acute, 0.22 ± 0.02 vs POST-Acute, 0.18 ± 0.02, V = 307, p = 1.47×10^−4^, Wilcoxon signed-rank test) but not on day-10 (PRE-Chronic, 0.27 ± 0.03 vs POST-Chronic, 0.28 ± 0.02, V = 145, p = 0.90, Wilcoxon signed-rank test). (F) Pre-stress information content (bits/spike) of place fields does not differ between day-1 and day-10 (PRE-Acute vs PRE-Chronic, p>0.05, KS-test). However, place cells active during RUN, before and after stress exposure, display a significant increase on day-1 (PRE-Acute, 1.96 ± 0.16 vs POST-Acute, 2.22 ± 0.14, V = 91, p = 0.002, Wilcoxon signed-rank test) but not on day-10 (PRE-Chronic, 1.68 ± 0.17 vs POST-Chronic, 1.59 ± 0.15, V = 167 p = 0.643, Wilcoxon signed-rank test). All box plots represent interquartile range (IQR, 25^th^-75^th^ percentiles), median is the thick line in the box and whiskers extend to 1.5 times the IQR. The black and red dotted lines on density plots display median values. * p<0.05, ** p < 0.01, *** p < 0.001, N = 5 mice.

### Differential impact of acute and chronic stress on spatial tuning of CA1 place cells

Our recordings from the dorsal CA1 region of the hippocampus (supplementary Fig. 1A) during baseline activity state (REST) yielded in a total of 180 pyramidal cells on day-1 (Acute) and 166 pyramidal cells on day-10 (Chronic). No major difference in firing rates was observed, although bursting activity showed a small, but significant, increase at the chronic time point (Table-1). Next, we assessed the impact of stress on mouse behaviour during track exploration (RUN) and observed no change between day-1 and day 10, as the total number of laps travelled by mice did not differ between sessions across days (2-way repeated measures ANOVA: main effect of day, F _(1, 4)_ = 0.564, p = 0.467; main effect of session, F _(1, 4)_ = 1.459, p = 0.250; interaction, F _(1, 4)_ = 0.001, p = 0.974). Similarly, stress did not disrupt the trial-averaged exploration speed on the track (2-way repeated measures ANOVA: main effect of day, F _(1, 4)_ = 3.106, p = 0.103; main effect of session, F _(1, 4)_ = 1.318, p = 0.273; interaction, F _(1, 4)_ = 0.08, p = 0.782). These data demonstrated that neither acute nor chronic stress strongly affected mouse locomotor behaviour.

**Table. 1.** Pyramidal cell properties during baseline activity state (SLP1) on day-1 and day-10 Impact of stress on CA1 pyramidal cells during baseline activity (REST) and exploration (RUN) *Top*; Comparisons of spiking and bursting properties of pyramidal cells during quite wakefulness/sleep (REST) between day-1 and day-10. *Bottom*; Comparisons of spiking and bursting properties of pyramidal “place” cells during pre-stress track exploration (RUN) between day-1 and day-10. Data is presented as Mean ± SEM, Wilcoxon rank-sum test, * p<0.05, ** p < 0.01, *** p < 0.001, N = 5 mice.

| **Parameters** | **First-day (REST)**<br>**(n=180 cells)** | **Last-day (REST)**<br>**(n=166 cell)** | **p-value** |
| --- | --- | --- | --- |
| Peak firing (Hz) | 4.60± 0.69 | 5.68± 0.59 | W=12556, *p = 0.01 |
| Mean firing (Hz) | 0.49± 0.07 | 0.53± 0.06 | W=13316, p = 0.08 |
| Complex Spike Index | 22.21± 0.95 | 23.48± 1.11 | W=14384, p = 0.55 |
| Burst duration | 7.72±0.19 | 7.96±0.22 | W=12189, *p = 0.0108 |
| Burst ratio | 0.32±0.01 | 0.36±0.01 | W=13306, p = 0.08 |
| Spikes per burst (n) | 2.25±0.01 | 2.32±0.02 | W=12267, ** p = 0.0051 |
| **Place cell properties** |  |  |  |
|  | **First-day (RUN)**<br>**(n=95 cells)** | **Last-day (RUN)**<br>**(n=101 cell)** | **p-value** |
| Peak firing (Hz) | 4.33± 0.30 | 4.94± 0.39 | W=4339, p = 0.248 |
| Mean firing (Hz) | 0.56± 0.06 | 0.76± 0.09 | W=4289, p = 0.201 |
| Complex Spike Index | 20.31± 1.16 | 21.39± 1.11 | W=4505, p = 0.463 |
| Burst duration | 7.40±0.10 | 7.23±0.16 | W=6824, p = 0.128 |
| Burst ratio | 0.34±0.01 | 0.36±0.01 | W=5870, p = 0.477 |
| Spikes per burst (n) | 2.26±0.02 | 2.27±0.02 | W=5869, p = 0.546 |

Next, we examined the impact of stress on the location-specific activation of CA1 pyramidal place cells during linear track exploration (O’Keefe and Dostrovsky, 1971). The fraction of neurons that passed place cell criteria during RUN (see Methods) was similar on the first and the last day of CIS (Fig. 1B) (day-1: 95/180 (53%) vs day-10: 101/166 (61%), p = 0.164, χ^2^ test), indicating that chronic stress did not alter activation of place cell ensembles. As expected, the mean firing of cells was significantly higher during RUN compared to the baseline REST session (Fig. 1C), with no discernible effect of repeated stress exposure (averaged firing; LMMs: main effect of day, F _(1, 538)_ = 3.988, p = 0.046; main effect of session, F _(1, 538)_ = 64.02, p = 7.708 × 10^−15^; interaction, F _(1, 538)_ = 0.018, p = 0.892). Thus, pyramidal cells increased their discharge rate during spatial coding and neither acute nor chronic stress affected this property of pyramidal cells.

We then performed detailed analysis of place cell properties. Place field size, defined as the proportion of the track that a place cell was active on, showed a main effect of day (Fig. 1D) (size; LMMs: main effect of day, F _(1, 369)_ = 8.660, p = 0.0035; main effect of session, F _(1, 369)_ = 1.201, p = 0.274; interaction, F _(1, 369)_ = 0.019, p = 0.890). Further, density distribution of field size during PRE-stress sessions changed after repeated stress such that compared to day-1, on day-10 a greater fraction of cells had larger place fields (KS-test, p = 0.049). Moreover, neurons that were active during both PRE and POST stress sessions displayed a decrease in field size after stress exposure on day-1 (PRE-Acute, 13.89 ± 1.46 vs POST-Acute, 11.25 ± 0.96, p= 0.009, Wilcoxon signed-rank test) but not on day-10 (PRE-Chronic, 19.71 ± 2.39 vs POST-Chronic, 17.81 ± 1.99, p = 0.316, Wilcoxon signed-rank test). Thus, place fields decreased in size after the acute stress, but expanded after repeated exposure to stress.

Altered place field size alone fails to capture all changes in spatial coding, as previous studies have reported that bigger place fields can be suggestive of both improved spatial coding (Hussaini et al., 2011) and a loss of spatial specificity (McHugh et al., 1996). Thus, we next assessed the impact of stress on spatial tuning by measuring the sparsity-index, a metric of spatial selectivity (Jung et al., 1994). The sparsity-index of individual place cells was also impacted by day (Fig. 1E) (sparsity; LMMs: main effect of day, F _(1, 369)_ = 8.931, p = 0.003; main effect of session, F _(1, 369)_ = 1.929, p = 0.166; interaction, F _(1, 369)_ = 2.017, p = 0.156). A further analysis of cells that were active before and after exposure to stress confirmed this result, as significantly lower sparsity-index was noticed after acute stress (PRE-Acute, 0.22 ± 0.02 vs POST-Acute, 0.18 ± 0.02, p = 1.47×10^−4^, Wilcoxon signed-rank test), but not after repeated stress (PRE-Chronic, 0.27 ± 0.03 vs POST-Chronic, 0.28 ± 0.02, p = 0.90, Wilcoxon signed-rank test). Further, spatial information content (bits/spike), a parameter which quantifies how much information about the mouse’s location is contained within the activity of a place cell (Skaggs et al., 1993), also showed effects of day (Fig. 1F) (information; LMMs: main effect of day, F _(1, 369)_ = 10.969, p = 0.001; main effect of session, F _(1, 369)_ = 2.586, p = 0.109; interaction, F _(1, 369)_ = 5.414, p = 0.0205). This was further confirmed as place cells active on the track before and after the exposure to stress also showed a significant increase in information content on day-1 (PRE-Acute, 1.96 ± 0.16 vs POST-Acute, 2.22 ± 0.14, p = 0.002, Wilcoxon signed-rank test) but not on day-10 (PRE-Chronic, 1.68 ± 0.17 vs POST-Chronic, 1.59 ± 0.15, p = 0.643, Wilcoxon signed-rank test) of the CIS protocol.

Overall, the smaller place fields and enhanced spatial tuning following the initial stress exposure suggest that acute stress improved hippocampal spatial coding.

### Differential impact of acute and chronic stress on exploration-associated theta and gamma oscillations

During exploratory behaviour the hippocampal local field potential (LFP) is dominated by prominent large-amplitude theta (6-12 Hz) oscillations (O’Keefe and Dostrovsky, 1971; Vanderwolf, 1969), which play a crucial role in the temporal organization of hippocampal activity (Buzsáki and Moser, 2013). Given the sharpened spatial tuning observed after acute stress, we next examined the impact of acute and chronic stress on theta oscillations. A comparison of the power spectral density of the LFP across sessions (Fig. 2B) revealed that theta oscillations were robustly present and power in the theta band was not affected by either acute or chronic stress, as no effect of day or session was observed (theta; 2-way repeated measures ANOVA: day, main effect of day F _(1, 19)_ = 2.7018, p=0.1262; main effect of session, F _(1, 19)_ = 0.0427, p = 0.8398; interaction, F _(1, 19)_ = 0.4823, p = 0.501).

**Fig. 2.**
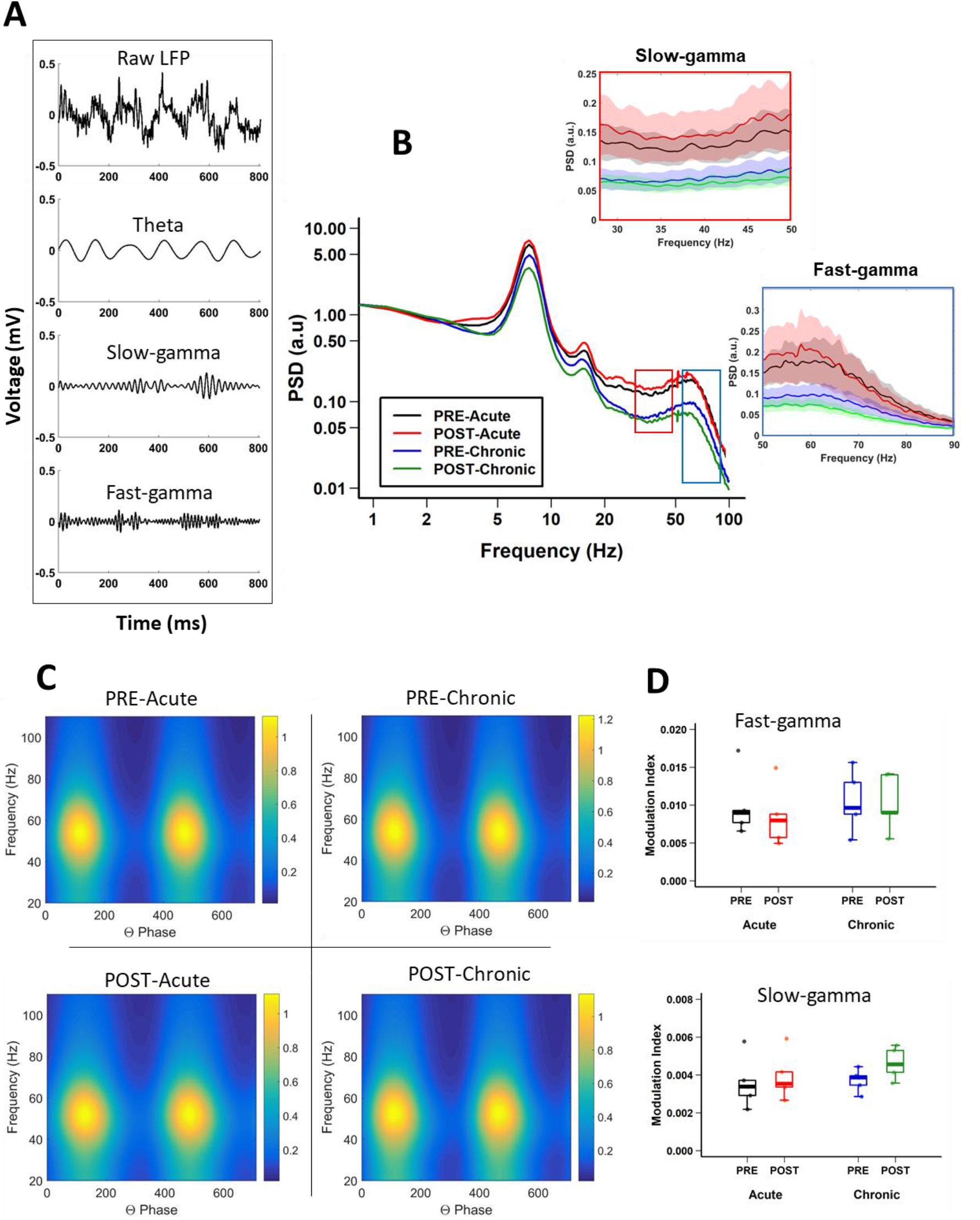
Impact of stress on CA1 oscillatory activity. (A) Representatives of unfiltered LFP during track exploration (top) and filtered (bottom), theta and slow-gamma and fast-gamma. (B) Power spectrum density (PSD) curves of CA1 local field potentials (LFPs) during linear track exploration (RUN) show no significant differences for theta (6-12 Hz) (Theta, 2-way repeated measure ANOVA: day, main effect of day F_(1, 19)_ = 2.7018, p=0.1262; main effect of session, F_(1, 19)_ = 0.0427, p =0.8398; interaction, F _(1, 19)_ = 0.4823, p = 0.501). Fast-gamma (55-90 Hz) showed an effect of day but not of session was noticed (FG, *right inset*; 2-way repeated measure ANOVA: main effect of day, F _(1, 19)_ = 6.7062, p=0.0237; main effect of session, F _(1, 19)_ = 0.1047, p =0.752 interaction, F _(1, 19)_ = 0.2074, p = 0.657). Slow-gamma (30-50 Hz) showed an effect of day but not of session (SG, *top inset* 2-way repeated measure ANOVA: main effect of day, F _(1, 19)_ = 8.3668, p=0.0135; main effect of session, F _(1, 19)_ = 0.0634, p = 0.805; interaction, F _(1, 19)_ = 0.3754, p = 0.551). (C) Representative examples of modulation of gamma amplitude by theta phase in dorsal CA1 pyramidal cell layer before (top) and after (bottom) stress exposure on day-1 (left) and day-10 (right) (D) Theta-FG phase-amplitude coupling (top) did not differ (2-way repeated measures ANOVA: day, main effect of day, F _(1, 4)_ = 0.839, p = 0.411; main effect of session, F _(1, 4)_ = 3.399, p = 0.139; interaction, F _(1, 4)_ = 4.953, p = 0.09). Similarly, theta-SG coupling (bottom) was not affected by either acute or chronic stress (2-way repeated measures ANOVA: day, main effect of day, F _(1, 4)_ = 0.592, p = 0.484; main effect of session, F _(1, 4)_ = 1.756, p = 0.256; interaction, F _(1, 4)_ = 1.02, p = 0.370). * p<0.05, ** p < 0.01, *** p < 0.001, N = 5 mice.

In addition to theta, the hippocampus displays occasional low-amplitude, high frequency gamma (30-90 Hz) oscillations (Bragin et al., 1995; Buzsáki et al., 2003; Colgin, 2016). Gamma oscillations consist of distinct subtypes with non-overlapping frequency ranges, slow (SG: 30-50 Hz) and fast (FG: 55-90 Hz) gamma (Alexander et al., 2018; Colgin, 2016; Middleton and McHugh, 2016), and enhanced gamma oscillatory activity has been suggested to reflect dynamic changes in excitatory input into CA1 (Buzsáki and Moser, 2013; Fries, 2015). Thus, we next assessed the impact of stress on these individual gamma bands. Significant decreases in fast-gamma power were evident on day-10 (Fig. 2B; FG; 2-way repeated measures ANOVA: main effect of day, F _(1, 19)_ = 6.7062, p=0.0237; main effect of session, F _(1, 19)_ = 0.1047, p = 0.752; interaction, F _(1, 19)_ = 0.2074, p = 0.657). Similarly, chronic stress also led to similar decreases in slow-gamma power (Fig. 2B; SG; 2-way repeated measures ANOVA: main effect of day, F _(1, 19)_ = 8.3668, p = 0.0135; main effect of session, F _(1, 19)_ = 0.0634, p = 0.805; interaction, F _(1, 19)_ = 0.3754, p = 0.551).

The amplitude of both fast and slow gamma oscillations has been shown to be modulated by the phase of slower underlying theta rhythm (Bragin et al., 1995; Canolty et al., 2006; Chrobak and Buzsáki, 1998) and this theta-phase gamma-amplitude coupling has been suggested to reflect local information processing in hippocampal circuits (Buzsáki and Wang, 2012; Tort et al., 2009). Thus, we next determined the impact of stress on theta-gamma coupling during periods when mice ran along the linear track linear (i.e. when prominent theta oscillations are known to be present) by calculating modulation index (MI), a measure of the strength of coupling between gamma-amplitude and theta phase (Tort et al., 2010). We found no changes in strength of either theta-fast gamma coupling (Fig. 2C, D) (theta-FG; 2-way repeated measures ANOVA: day, main effect of day, F _(1, 4)_ = 0.839, p = 0.411; main effect of session, F _(1, 4)_ = 3.399, p = 0.139; interaction, F _(1, 4)_ = 4.953, p = 0.09) nor theta-slow gamma coupling (theta-SG; 2-way repeated measures ANOVA: day, main effect of day, F _(1, 4)_ = 0.592, p = 0.484; main effect of session, F _(1, 4)_ = 1.756, p = 0.256; interaction, F _(1, 4)_ = 1.02, p = 0.370).

Thus, LFP power analysis indicated that while the first exposure to stress did not alter theta and gamma oscillatory activity, repeated stress led to suppression of SG and FG power, but had no impact on cross-frequency coupling between theta and gamma.

### Impact of acute and chronic stress on temporal coding (LFP-spike interaction)

In addition to rate coding (location-specific spiking), place cells also display temporal coding, reflecting in their preference for spiking at specific phases of the concurrent oscillations (Csicsvari et al., 1999; Fox et al., 1986; O’Keefe, 1976). It has been hypothesized that temporal coding supports transient activation of place cell ensembles, a phenomena central to spatial information processing (Buzsáki, 2010; Harris et al., 2003; Lever et al., 2014; O’Keefe and Burgess, 2005). Knowing that acute and chronic stress differentially alter the place cell rate code, we next asked if they differentially impact temporal coding by assessing the strength and phase preference of CA1 place cell spiking to theta and gamma oscillations. Similar to a previous report (Jones and Wilson, 2005), the majority (66~73%) of CA1 place cells demonstrated significant modulation by theta (Rayleigh test of uniformity p < 0.05) and neither acute nor chronic stress affected this distribution (Table-2; p = 0.814, χ^2^ test). Further, as expected based on earlier studies (Csicsvari et al., 1999; Jadhav et al., 2016; Jones and Wilson, 2005), the majority of neurons displayed a preference to spike near the trough of the theta oscillation (Fig. 3A) and this mean preferred phase for theta-modulated cells was not affected by stress (supplementary Fig. 1B; phase; Circular ANOVA, F _(3, 155)_ = 1.305, p = 0.274). Interestingly however, the strength of theta-phase locking (Fig. 3B) was significantly increased specifically after acute stress (MI; 2-way ANOVA: main effect of day, F _(1, 155)_ = 5.296, p = 0.0023; main effect of session, F _(1, 155)_ = 3.009, p = 0.085; interaction, F _(1, 155)_ = 6.579, p = 0.011); post hoc Tukey’s test, PRE-Acute vs PRE-Stress, p = 0.013, POST-Acute vs POST-Chronic, p = 0.004).

**Table 2.** Distribution of place cells phase-locked to theta and gamma oscillation on day-1 and day-10.

| **Rhythm** | **PRE-Acute** | **POST-Acute** | **PRE-Chronic** | **PRE-Chronic** | **Statistics** |
| --- | --- | --- | --- | --- | --- |
| Theta | 35/48 (73%) | 26/38 (68%) | 53/73 (73%) | 45/68 (66%) | p = 0.814, $\chi^2$ test |
| Fast-gamma | 30/66 (45%) | 26/63 (41%) | 34/78 (44%) | 30/74 (40%) | p = 0.935, $\chi^2$ test |
| Slow-gamma | 27/74 (36%) | 30/87 (34%) | 29/91 (32%) | 31/87 (36%) | p = 0.928, $\chi^2$ test |

**Fig. 3.**
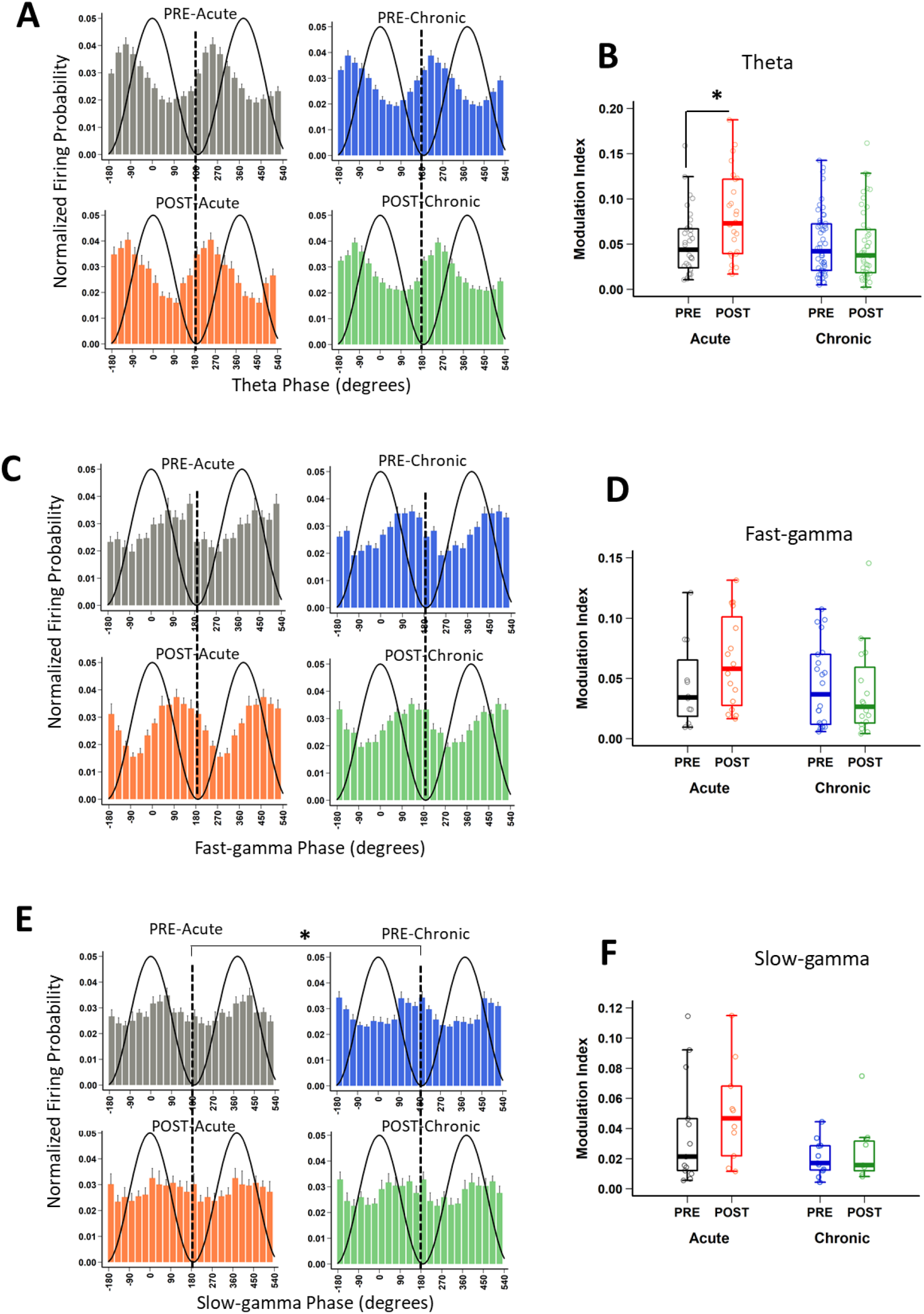
Impact of stress on phase-locking properties of CA1 place cells to theta and gamma oscillations. (A) The spiking probability plotted as a function of the phase of theta for significantly theta-modulated place cell populations (Rayleigh p<0.05). Population spiking probability is elevated around the trough and ascending phase of theta (0/360° set for theta peak, 180° for theta trough). (B) The strength of theta-phase locking (MI) is altered by stress (2-way ANOVA: main effect of day, F _(1,155)_ = 5.296, p = 0.0023; main effect of session, F _(1, 155)_ = 3.009, p = 0.085; interaction, F_(1, 155)_ = 6.579, p = 0.011); post hoc Tukey’s test, PRE-Acute vs PRE-Stress, p=0.013, Post-Acute vs Post-Chronic, p=0.004). (C) The spiking probability plotted as a function of the phase of fast-gamma (FG) for significantly FG-modulated place cell populations (Rayleigh p<0.05). Population spiking probability is elevated around the trough and descending phase of FG (0/360° set for FG peak, 180° for FG trough) but stress did not affect mean phase: (phase, Circular ANOVA, F _(3,62)_ = 0.919, p=0.437). (D) The strength of FG-phase locking (MI) remains unaltered by stress (FG; 2-way ANOVA: main effect of day, F_(1,62)_ = 1.923, p = 0.171; main effect of session, F_(1,62)_ = 0.348, p = 0.558; interaction, F_(1,62)_ = 1.808, p = 0.184). (E) The spiking probability is elevated around the descending phase of slow-gamma (0/360° set for SG peak, 180° for SG trough) and phase was altered by chronic stress (phase; Circular ANOVA, F _(3,37)_ = 5.057, p=0.005; post hoc Watson-Wheeler test, PRE-Acute vs PRE-Chronic, p=0.036. (F) The strength of SG-phase locking (MI) was altered by stress (SG; 2-way ANOVA: main effect of day, F_(1,37)_ = 4.856, p = 0.034; main effect of session, F_(1,37)_ = 1.101, p = 0.300; interaction, F_(1,37)_ = 0.123, p = 0.727). Boxplots represent interquartile range (IQR, 25^th^-75^th^ percentiles), median is the thick line in the box and whiskers extend to 1.5 times the IQR., * p<0.05, ** p < 0.01, *** p < 0.001, N = 5 mice.

Similar to the modulation of spiking by theta, the precise timing of pyramidal cell firing can also be entrained by gamma oscillations (Csicsvari et al., 2003). Thus, we next performed the spike phase-locking analysis of gamma oscillations during high velocity periods (speed > 6cm/s) on the track. A large fraction (40~45%; Rayleigh test of uniformity p < 0.05) of CA1 place cell population displayed a significant phase preference during FG and neither acute nor chronic stress affected this distribution (Table-2; p = 0.935, χ^2^ test). Moreover, stress did not alter the preferred phase (phase; Circular ANOVA, F _(3,108)_ = 1.932, p=0.129; supplementary Fig. 1C) or strength (MI; 2-way ANOVA: main effect of day, F (1,108) = 1.736, p = 0.190; main effect of session, F _(1, 108)_ = 1.095, p = 0.298; interaction, F _(1, 108)_ = 0.149, p = 0.701) of the phase locking of CA1 pyramidal cells. As gamma oscillations are more transient than theta during locomotion, we next focused on periods of strong FG on the track regardless of animal’s speed or position on the track (see Methods) and again observed that stress did not alter either the preferred phase (Fig. 3C; phase: Circular ANOVA, F _(3,62)_ = 0.919, p=0.437) or the strength of fast-gamma-phase locking (Fig. 3D; MI: 2-way ANOVA: main effect of day, F _(1,62)_ = 1.923, p = 0.171; main effect of session, F _(1, 62)_ = 0.348, p = 0.558; interaction, F _(1, 62)_ = 1.808, p = 0.184).

Unlike FG, the preferred phase of the cells modulated by SG was more variable across the population (supplementary Fig. 1 D) during high velocity periods on the track. The proportion of CA1 place cells with a significant SG phase preference was ~ 32-36%; (Rayleigh test of uniformity p < 0.05) and showed no differences across four sessions (Table-2; p = 0.928, χ^2^ test). Nonetheless, following chronic stress we did observe a small, yet significant change in the mean preferred phase (supplementary Fig. 1 D; phase: Circular ANOVA, F _(3,113)_ = 6.862, p=2.75 × 10^−4^), but noticed no change in the strength of modulation (MI: 2-way ANOVA: main effect of day, F _(1,113)_ = 0.182, p = 0.671; main effect of session, F_(1, 113)_ = 0.026, p = 0.872; interaction, F_(1, 113)_ = 2.463, p = 0.119). Finally, when we focused on the phase-locking of place cells specifically during periods of strong SG regardless of animal’s speed or position on the track, we found that chronic stress again altered both the preferred phase (Fig. 3E-phase; Circular ANOVA, F _(3,37)_ = 5.057, p=0.005; post hoc Watson-Wheeler test, PRE-Acute vs PRE-Chronic, p=0.036) and the strength of SG-phase locking (Fig. 3F; MI: 2-way ANOVA: main effect of day, F _(1,37)_ = 4.856, p = 0.034; main effect of session, F_(1, 37)_ = 1.101, p = 0.300; interaction, F_(1, 37)_ = 0.123, p = 0.727).

Thus, while first exposure to stress increased the strength of theta-phase locking demonstrating the facilitatory effects of acute stress, on temporal coding, chronic stress disrupted temporal coding as the mean phase and the strength of phase-locking of place cells to slow-gamma oscillations was altered on day-10.

## Discussion

Despite reports that acute stress positively impacts cognition, including hippocampal information processing (Henckens et al., 2009; Kirby et al., 2013; Yuen et al., 2009), it is not yet clear how this is reflected in hippocampal place cell activity and LFP-spike interactions, two neural processes supporting spatial coding (Buzsáki, 2010; Lever et al., 2014; O’Keefe and Dostrovsky, 1971). Here, we show that after the first exposure to stress (Acute stress) CA1 place cells displayed refined spatial coding (Fig. 1D), increased information content (Fig. 1F) and decreased sparsity-index (Fig. 1E). Further, chronic, but not acute stress, led to decreased LFP power in the slow-gamma (SG; 30-50 Hz) and fast-gamma (FG 55-90 Hz) bands (Fig. 2 B) and an increase in place field size. Furthermore, the strength of theta-phase locking to CA1 place cells increased after acute stress (Fig. 3A, B), however the mean phase of slow-gamma phase-locking was altered after stress became chronic (Fig. 3E). Together, these results indicate that acute stress has a facilitatory impact on hippocampal information coding, while chronic stress impairs it.

Stress impacts on hippocampal functionality have been hypothesized to follow a U-shaped curve, where exposure to acute stress facilitates, while chronic stress disrupts, hippocampal function (McEwen et al., 2016; Salehi et al., 2010). Our results of enhanced information content in place fields and increased strength of phase locking after acute stress, as well as broader place fields and suppressed gamma power after repeated stress are consistent with stress exerting a U-shaped impact on hippocampal function in the intact brain. Rate and temporal coding of CA1 pyramidal cells aid spatial information processing (O’Keefe, 1976; O’Keefe and Burgess, 2005; O’Keefe and Recce, 1993). Since acute stress facilitated both type of coding (i.e., improved spatial tuning and strength of theta-phase locking), it is not farfetched to assume that acute stress effects on hippocampal spatial coding are indeed facilitatory in nature. Mechanistically, the facilitatory effects of acute stress on hippocampal coding are likely brought about by the combined action of a cocktail of neuromodulators released by stress-induced activation of sympatho-adrenal medullary (SAM)-pathways (Cadle and Zoladz, 2015; Gunn and Baram, 2017). Future studies are needed to further investigate the role of SAM-activated neuromodulation on CA1 spatial coding.

Instantaneous coupling between theta and gamma oscillations in hippocampal networks is thought to represent dynamic processing in hippocampal circuits (Buzsáki and Wang, 2012). A previous CIS study using evoked auditory potentials also noted a decrease in gamma power following chronic stress and concluded poor functional connectivity within the hippocampal circuitry (Ghosh et al., 2013). The same conclusion was also reached by Passecker and colleagues (2011) who studied the impact of repeated exposure to photic stress on hippocampal spatial coding. Gamma oscillations route information flow in hippocampal circuits such that slow CA1 gamma reflects interactions between CA3-CA1 neuronal networks (Montgomery and Buzsáki, 2007), while fast CA1 gamma indicates CA1-MEC interactions (Colgin, 2016; Colgin et al., 2009). Our observation of decreased slow and fast gamma power following chronic, but not acute, stress exposure, may directly reflect the poor functional connectivity and information transfer in hippocampal-entorhinal circuits in chronically stressed subjects. Importantly, functional connectivity was not altered after acute stress, as place maps were more informative of animal’s location in space.

What may lead to weakened functional connectivity in hippocampal circuits in response to repeated stress? Earlier studies have reported that chronic, but not acute stress, causes dendritic shortening and debranching and synaptic loss on apical branches of pyramidal cells in areas CA3 and CA1 (Conrad et al., 1999; Magariños and McEwen, 1995; Sandi et al., 2003; Sousa et al., 2000). Hitherto, the functional consequences of these structural changes have not been well understood. Since apical dendritic branches of CA1 pyramidal cells are the loci of Schaffer collateral inputs (from CA3) and temporoammonic pathways from medial entorhinal (MEC) cortex, chronic stress-induced CA1 dendritic shrinkage likely reflects poor information flow into CA1 circuits (Colgin et al., 2009). Knowing that CA1 SG oscillations reflect interactions between CA1 and CA3/CA2 circuitry (Alexander et al., 2018; Colgin et al., 2009; Middleton and McHugh, 2016), while FG occurs during interactions between area CA1 and medial entorhinal cortical circuits (Colgin et al., 2009; Kemere et al., 2013), it is not surprising that chronic (but not acute stress) causes a decrease in SG and FG power. In addition, altered AMPA-dependent synaptic plasticity can also alter inhibitory-excitatory balance and contributes to gamma phase-locking of CA1 pyramidal cells (Kitanishi et al., 2015). Knowing that chronic stress alters hippocampal synaptic plasticity (Alfarez et al., 2003) and AMPA-dependent synaptic transmission in temporoammonic-CA1 pathway (Kallarackal et al., 2013), it is likely that chronic stress-induced altered synaptic plasticity is another potential candidate underlying chronic stress phenotypes noticed in this study. Further, inhibitory neuronal activity plays a key role in the generation of gamma oscillations, as well as the phase-locking of pyramidal cells to gamma oscillations (Bartos et al., 2002; Buzsáki and Wang, 2012). Reports that chronic stress causes decreases in hippocampal PV^+^ inhibitory neuronal density by ~ 20-25% (Csabai et al., 2017; Zaletel et al., 2016) suggests that chronic stress-induced loss of inhibitory neurons may also contributed to decreased gamma power as well as altered gamma phase-locking by place cells in chronically stressed subjects. Future studies will have to assess the differential contributions of chronic-stress-induced altered inhibition, synaptic plasticity and apical dendritic atrophy of pyramidal cells in altered place cell activity, gamma oscillations and phase-locking phenotypes observed in this study.

A decrease in gamma (30-90 Hz) power and broadening of place field size after repeated stress exposure indicates that acute and chronic stress differentially alter information coding in the CA1 subregion. In view of reports that hippocampal phase-locking is altered in neurodegenerative disease models (Booth et al., 2016; Mably et al., 2017), for which stress is a risk factor (Bisht et al., 2018), it is not surprising that chronic stress altered some phase-locking parameters. These data further add to an accumulating evidence that repeated stress negatively impacts spatial coding (Chattarji et al., 2015; Kim et al., 2007; Tomar et al., 2015). Spike-oscillation interactions are responsible for not only local computations within a circuit but also coordinate activity across distant but connected circuits (Buzsáki and Freeman, 2015; Colgin, 2016; Harris and Gordon, 2015; Makino et al., 2019; Shin and Jadhav, 2016). Thus, our results of altered oscillatory and place cell activity have implication for neural computations across various memory-related circuits connected to the hippocampus.

In conclusion, our results of acute-stress induced increased information content of place cells and strengthening of phase-locking to theta oscillations further support the idea that acute stress facilitates hippocampal neural computations. Based on these findings, we propose that acute and chronic stress differentially, likely opposingly, influence hippocampal information processing.

## Acknowledgements

We thank all members of the CBP laboratory for their support, the Advanced Manufacturing Support Team, RIKEN Center for Advanced Photonics for their assistance in microdrive production and Lalitha Krishnan for assistance with figure generation. This works was supported by Grant-in-Aid for Scientific Research from MEXT (19H05646; T.J.M), Grant-in-Aid for Scientific Research on Innovative Areas from MEXT (19H05233; T.J.M) and RIKEN BSI and CBS (T.J.M).

## Author contributions

A.T. and T.J.M conceived the study. A.T. performed all experiments. A.T. D.P. and T.J.M. analysed the data. A.T., and T.J.M. wrote the manuscript with inputs from D.P. Funding provided by T.J.M.

## Correspondence and request for material

Any request for data or code should be addressed to A.T. or T.J.M.

## Material and Methods

### Animals

All experiments were performed using male C57BL/6J mice. A total of 5 mice, aged between 3 and 6 months, were used for this study. The data related to the physiology during stress of these mice was previously reported in (Tomar et al., 2021). Mice were maintained on a 12-h light-dark cycle with *ad libitum* access to food and water. All procedures were approved by the RIKEN Institutional Animal Care and Use Committee and complied with the National Institutes of Health guide for the care and use of Laboratory animals (NIH Publications No. 8023, revised 1978). All efforts were made to minimise animal suffering and to reduce the number of animals used.

### Experimental design and stress protocol

Mice were habituated to the small sleep-box as well as a linear track daily and after surgery mice were again habituated to sleep-box in which later all “rest” data was collected. Thus, mice were completely habituated to the experimenter, room, sleep box, etc., minimizing the contribution of other (non-stress) repetitive factors/experiences to the changes we observed in the physiology of the hippocampus. Mice underwent the same chronic immobilization stress (CIS) protocol as described previously (Tomar et al., 2015). Briefly, mice experienced complete immobilization (2h/d for 10 consecutive days: Fig. 1A) in rodent immobilization bags, without access to either food or water. During the actual experiment, all mice experienced a familiar track twice, the first before (PRE) and second after the stress exposure (POST), on the first day (Acute) and the last day (Chronic) of a chronic immobilization stress (CIS) paradigm thus providing us with four conditions, i) PRE-Acute, ii) POST-Acute, iii) PRE-Chronic iv) POST-Chronic. Each track (RUN) epoch was bracketed by Rest-state (REST) epoch and each epoch was ~ 30 min.

### Surgery, recording and histology

Mice were anaesthetized using Avertin (2, 2, 2-tribromoethanol; Sigma-Aldrich, 476 mg/kg, i.p.) and were surgically implanted with a microdrive (manufactured with the assistance of the Advanced Manufacturing Support Team, RIKEN Center for Advanced Photonics, Japan). The microdrive housed eight independently movable tetrodes (14 μm diameter, nichrome) and was placed above right dorsal hippocampus (coordinates from bregma: AP −1.8 mm; ML + (1.2 mm). Prior to surgery, tetrodes were gold plated to lower impedance down to a range of 100–250 kΩ. Tetrodes were gradually lowered over the course of several days, such that by the start of the experiment they reached the CA1 stratum pyramidale. Data were acquired using a 32-channel Digital Lynx 4S acquisition system (Neuralynx, Bozeman, MT). Signals were sampled at 32,556 Hz and spike waveforms were filtered between 600 Hz and 6 kHz. Skull screws located above the cerebellum served as a ground, and a tetrode positioned in the superficial layers of the neocortex, and devoid of spiking activity, was used for referencing. 3-4 weeks after surgery, when all tetrodes reached the CA1 stratum pyramidale, evident by multiple large amplitude spikes and SPW-Rs, the experiment was initiated. During REST epochs the mice were located in a small circular sleep box (15 cm diameter). At the conclusion of the experiment mice underwent terminal anaesthesia (Avertin), and electric current (30 μA, for 8 s) was administered through each electrode to mark their locations. Transcardial perfusion was carried by using saline followed by 4% paraformaldehyde (PFA) followed by a further 24 h fixation in 4% PFA. Brains were sliced using a vibratome (Leica) to prepare coronal slices (50 μm thick) and inspected by standard light microscopy to confirm electrode placement.

### Unit isolation and spike analysis

Spike sorting was performed by automatic spike sorting program (KlustaKwik; (Harris et al., 2000)), followed by manual adjustments of the cluster boundaries using SpikeSort3D software (Neuralynx). Candidate clusters with < 0.5% of spikes displaying an inter-spike-interval shorter than 2 ms, a total number of spikes exceeding 50, having a cluster isolation distance value (Schmitzer-Torbert et al., 2005) ≥ 10, spike halfwidth (peak-to-trough) >170 μs and complex spike index (CSI) (McHugh et al., 1996) >5 were considered as pyramidal cells and were used for further analysis.

### Place cell properties

Pyramidal cells that were active during period of exploration (RUN) with a speed >2cm/s on the linear track, had field size ≥ 6 bins and covered less than 50% of the track were considered place cells. Peak firing rate was defined as the firing rate in the spatial bin containing the maximal value within each firing rate map. Place field size was defined as the number of spatial bins where place cell field firing exceeded 20% of the peak firing rate. Mean firing rate was calculated by dividing number of spikes which occurred within periods when velocity exceeded 2 cm/sec by that period’s duration and then these values were averaged. Complex Spike Index (CSI) is defined as CSI = 100 * (pos - neg), where ‘pos’ is the number of inter-spike intervals positively contributing to CSI, that is, preceding spikes with larger amplitudes and following spikes with smaller amplitudes (complex bursts) occurring within 3 ms (refractory period) and 15 ms (maximum inter-spike interval defining a burst); ‘neg’ is the number of inter-spike intervals that contribute negatively to CSI, i.e. violating either or both these rules. A “burst” was defined as at least two spikes occurring within 10 ms time bin. The burst detection and analysis was performed using Matlab scripts previously described in (Bakkum et al., 2014). Place field ‘sparsity’ was computed as previously described in (Resnik et al., 2012). Briefly, the ‘sparsity’ was defined as a number ranging from 0 to 1, were 0 corresponding to a firing rate map which consists of equal firing rate values in every visited spatial bin. Firing rate map with sparsity value 1 corresponds to the case when all the spikes generated by any given cell occurred in a single spatial bin. Spatial Information (SI, bits/spike) was calculated as previously reported (Skaggs et al., 1993); Briefly SI = sum(P_spk_(i) * log2(P_spk_(i) / P_occ_(i)), where P_spk_(i) is the probability of spiking in bin ‘i’ and ‘P_occ_(i)’ is the occupancy probability in bin ‘i’. The ‘P_spk_’ and ‘P_occ_’ values were computed from the rate and occupancy maps respectively.

### Power Spectral Density

The Power Spectral Density (PSD) during exploratory behaviour was calculated by using Welch’s averaged modified periodogram method with 2048 samples (1.26 s) window size, 50% overlap and 4096 FFT points (2.52 s) resulting a time-varying spectrogram. The PSD curves corresponding to time bins when animal’s velocity was above 6 cm/sec were averaged yielding single PSD curve for each of 4 experimental conditions. In order to account for power fluctuations caused by differences in position/impedance of the electrodes and make PSD values comparable across mice, we normalized each PSD curve by its own mean power within (0–3 Hz) band.

### Theta/Gamma Phase-locking to spikes

The phase relationship between spikes and theta LFP was calculated as previously described (Siapas et al., 2005). Briefly, instantaneous theta phase was derived from the Hilbert-transformed LFP trace filtered in the theta band (6–12 Hz). Peaks and troughs were assigned 0- and 180-degree phases respectively, with spike phase calculated using interpolation, a method not sensitive to theta wave asymmetry. The resultant phases were converted to firing probability histograms (10 degree bin size), while limiting spikes to time periods when animal’s velocity exceeded 6 cm/sec. Significance of the phase locking, preferred firing phase, strength of modulation and statistical comparison of phase values were calculated using functions from Circular Statistics Toolbox (Berens, 2009). Gamma/spikes modulation was computed in a similar manner; the calculation was performed using LFP traces filtered in slow gamma (30-50 Hz) and fast gamma (55-90 Hz) frequency bands. Due to transient nature of gamma oscillations, additional gamma ‘bursts’ detection was performed by calculating time periods when instantaneous power (absolute value of Hilbert transform) of gamma-band filtered LFP trace exceeded various threshold values (in Standard Deviations, 0.5SD, 1SD, 2SD) above mean value of the trace.

### Cross-Frequency Coupling between theta and gamma oscillations

Cross-Frequency Coupling (CFC) was calculated as described previously (Tort et al., 2010). To reliably detect the phenomena, relatively long chunks of LFP representing consistent behaviour state are necessary, thus time periods when the mouse was running along the track were used in this analysis. LFP data of each lap was first down sampled to 800 Hz, z-scored and converted to time varying power over multiple frequency bands matrices by using wavelet transform (mother wavelet function: Morlet, wavelet parameter: 5). Then modulation index (MI) values were calculated for each pair of low (4-20 Hz) and high (30-300 Hz) frequency bands.

Significance of the MI values was assessed by using permutation method (N_perm_ = 200), for details see (Tort et al., 2010).

### Statistical analysis

All statistical analyses were performed in R software (3.3.2). The normality of distributions was not assumed, so comparisons were made using non-parametric statistics. For between-group comparison Wilcoxon rank-sum tests was used, while for cells matched between two epochs, Wilcoxon signed-rank tests were used to test the equality of medians. For two-way ANOVAs (*aov* function, stats package) followed by Tukey’s honestly significant difference (HSD) test. (*TukeyHSD* function, stats package). Overall differences in place cell properties were assessed using linear mixed effects models (LMMs), where mouse identity was specified as a random factor and day and behaviour state were specified as fixed factors. The output of the *lmer* function was summarized as an ANOVA table (*anova* function, stats package). Similarly, comparisons for power distributions across various frequency bands in LFP signals were assessed using LMMs, where mouse identity was specified as a random factor and frequency bands as categorical variables were specified as fixed factors. Correlation between parameters was calculated using Pearson’s correlation coefficient analysis (base package). Dependence of a parameter on another was calculated by employing standardized major axis (SMA) regression (*sma* function, smatr package). Comparisons between regression lines were made by likelihood ratio tests (*sma* function, smatr package). For density curve analysis, the Kolmogorov-Smirnov test was employed (*ks.test,* stats package). For phase-locking analysis, statistical analyses were performed on 10 degree binned data however for visualization purposes data is presented in 30 degree bins. Boxplots represent Interquartile Range (IQR, 25^th^-75^th^ percentiles), median is the thick line housed in the box and whiskers extend to 1.5 times the IQR. No data points were removed as outliers either for making boxplots or for statistical analysis, however for visualization purposes, axes of graphs were readjusted. All statistical tests used were two-sided, and unless otherwise stated, the significance threshold for all tests was set at p < 0.05 and p-values are shown as follows: * p < 0.05; ** p < 0.01; *** p < 0.001.

## Data and code availability

The data and MATLAB codes that support the findings of this study are available from corresponding authors upon request.

---

**Supplementary Fig. 1.**
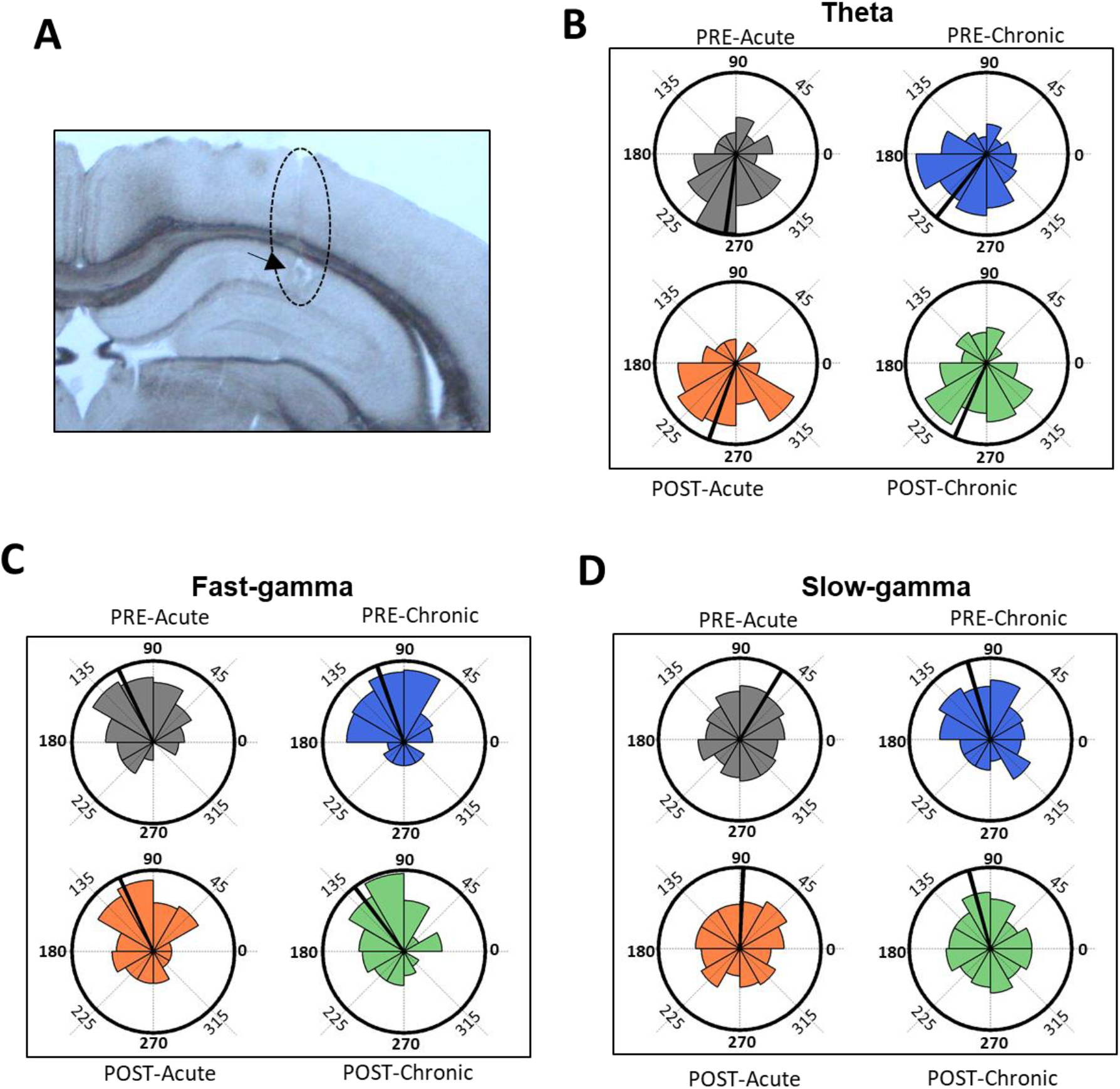
Impact of stress on phase-locking properties of CA1 place cells to theta oscillations. (A) Coronal section of the hippocampus showing the tetrode locations (black arrows) at the CA1 pyramidal layer. (B) Circular histograms display preferred phase of all place cells during theta (B), FG (C) and SG (D). The thick line in each circular histogram depicts averaged phase across all cells.

## References

Alexander, G.M., Brown, L.Y., Farris, S., Lustberg, D., Pantazis, C., Gloss, B., Plummer, N.W., Jensen, P., and Dudek, S.M. (2018). CA2 neuronal activity controls hippocampal low gamma and ripple oscillations. ELife 7, e38052.

Alfarez, D.N., Joëls, M., and Krugers, H.J. (2003). Chronic unpredictable stress impairs long-term potentiation in rat hippocampal CA1 area and dentate gyrus in vitro. Eur. J. Neurosci. 17, 1928–1934.

Bakkum, D.J., Radivojevic, M., Frey, U., Franke, F., Hierlemann, A., and Takahashi, H. (2014). Parameters for burst detection. Front. Comput. Neurosci. 7.

Bartos, M., Vida, I., Frotscher, M., Meyer, A., Monyer, H., Geiger, J.R.P., and Jonas, P. (2002). Fast synaptic inhibition promotes synchronized gamma oscillations in hippocampal interneuron networks. Proc. Natl. Acad. Sci. 99, 13222–13227.

Berens, P. (2009). CircStat: A MATLAB Toolbox for Circular Statistics. J. Stat. Softw. 31, 1–21.

Bisht, K., Sharma, K., and Tremblay, M.-È. (2018). Chronic stress as a risk factor for Alzheimer’s disease: Roles of microglia-mediated synaptic remodeling, inflammation, and oxidative stress. Neurobiol. Stress 9, 9–21.

Booth, C.A., Witton, J., Nowacki, J., Tsaneva-Atanasova, K., Jones, M.W., Randall, A.D., and Brown, J.T. (2016). Altered Intrinsic Pyramidal Neuron Properties and Pathway-Specific Synaptic Dysfunction Underlie Aberrant Hippocampal Network Function in a Mouse Model of Tauopathy. J. Neurosci. 36, 350–363.

Bragin, A., Jandó, G., Nádasdy, Z., Hetke, J., Wise, K., and Buzsáki, G. (1995). Gamma (40-100 Hz) oscillation in the hippocampus of the behaving rat. J. Neurosci. Off. J. Soc. Neurosci. 15, 47–60.

Buzsáki, G. (2010). Neural syntax: cell assemblies, synapsembles, and readers. Neuron 68, 362–385.

Buzsáki, G., and Freeman, W. (2015). Editorial overview: brain rhythms and dynamic coordination. Curr. Opin. Neurobiol. 31, v–ix.

Buzsáki, G., and Moser, E.I. (2013). Memory, navigation and theta rhythm in the hippocampal-entorhinal system. Nat. Neurosci. 16, 130–138.

Buzsáki, G., and Wang, X.-J. (2012). Mechanisms of gamma oscillations. Annu. Rev. Neurosci. 35, 203–225.

Buzsáki, G., Buhl, D.L., Harris, K.D., Csicsvari, J., Czéh, B., and Morozov, A. (2003). Hippocampal network patterns of activity in the mouse. Neuroscience 116, 201–211.

Cadle, C.E., and Zoladz, P.R. (2015). Stress time-dependently influences the acquisition and retrieval of unrelated information by producing a memory of its own. Front. Psychol. 6.

Canolty, R.T., Edwards, E., Dalal, S.S., Soltani, M., Nagarajan, S.S., Kirsch, H.E., Berger, M.S., Barbaro, N.M., and Knight, R.T. (2006). High Gamma Power Is Phase-Locked to Theta Oscillations in Human Neocortex. Science 313, 1626–1628.

Chattarji, S., Tomar, A., Suvrathan, A., Ghosh, S., and Rahman, M.M. (2015). Neighborhood matters: divergent patterns of stress-induced plasticity across the brain. Nat. Neurosci. 18, 1364–1375.

Chrobak, J.J., and Buzsáki, G. (1998). Gamma Oscillations in the Entorhinal Cortex of the Freely Behaving Rat. J. Neurosci. 18, 388–398.

Colgin, L.L. (2016). Rhythms of the hippocampal network. Nat. Rev. Neurosci. 17, 239–249.

Colgin, L.L., Denninger, T., Fyhn, M., Hafting, T., Bonnevie, T., Jensen, O., Moser, M.-B., and Moser, E.I. (2009). Frequency of gamma oscillations routes flow of information in the hippocampus. Nature 462, 353–357.

Conrad, C.D., LeDoux, J.E., Magariños, A.M., and McEwen, B.S. (1999). Repeated restraint stress facilitates fear conditioning independently of causing hippocampal CA3 dendritic atrophy. Behav. Neurosci. 113, 902–913.

Csabai, D., Seress, L., Varga, Z., Ábrahám, H., Miseta, A., Wiborg, O., and Czéh, B. (2017). Electron Microscopic Analysis of Hippocampal Axo-Somatic Synapses in a Chronic Stress Model for Depression. Hippocampus 27, 17–27.

Csicsvari, J., Hirase, H., Czurkó, A., Mamiya, A., and Buzsáki, G. (1999). Oscillatory coupling of hippocampal pyramidal cells and interneurons in the behaving Rat. J. Neurosci. Off. J. Soc. Neurosci. 19, 274–287.

Csicsvari, J., Jamieson, B., Wise, K.D., and Buzsáki, G. (2003). Mechanisms of gamma oscillations in the hippocampus of the behaving rat. Neuron 37, 311–322.

Fox, S.E., Wolfson, S., and Ranck, J.B. Jr, (1986). Hippocampal theta rhythm and the firing of neurons in walking and urethane anesthetized rats. Exp. Brain Res. Exp. Hirnforsch. Expérimentation Cérébrale 62, 495–508.

Fries, P. (2015). Rhythms for Cognition: Communication through Coherence. Neuron 88, 220–235.

Ghosh, S., Laxmi, T.R., and Chattarji, S. (2013). Functional connectivity from the amygdala to the hippocampus grows stronger after stress. J. Neurosci. Off. J. Soc. Neurosci. 33, 7234–7244.

Goutagny, R., Gu, N., Cavanagh, C., Jackson, J., Chabot, J.-G., Quirion, R., Krantic, S., and Williams, S. (2013). Alterations in hippocampal network oscillations and theta-gamma coupling arise before Aβ overproduction in a mouse model of Alzheimer’s disease. Eur. J. Neurosci. 37, 1896–1902.

Gunn, B.G., and Baram, T.Z. (2017). Stress and Seizures: Space, Time and Hippocampal Circuits. Trends Neurosci. 40, 667–679.

Harris, A.Z., and Gordon, J.A. (2015). Long-range neural synchrony in behavior. Annu. Rev. Neurosci. 38, 171–194.

Harris, K.D., Henze, D.A., Csicsvari, J., Hirase, H., and Buzsáki, G. (2000). Accuracy of Tetrode Spike Separation as Determined by Simultaneous Intracellular and Extracellular Measurements. J. Neurophysiol. 84, 401–414.

Harris, K.D., Csicsvari, J., Hirase, H., Dragoi, G., and Buzsáki, G. (2003). Organization of cell assemblies in the hippocampus. Nature 424, 552–556.

Henckens, M.J.A.G., Hermans, E.J., Pu, Z., Joëls, M., and Fernández, G. (2009). Stressed Memories: How Acute Stress Affects Memory Formation in Humans. J. Neurosci. 29, 10111–10119.

Hussaini, S.A., Kempadoo, K.A., Thuault, S.J., Siegelbaum, S.A., and Kandel, E.R. (2011). Increased size and stability of CA1 and CA3 place fields in HCN1 knockout mice. Neuron 72, 643–653.

Jadhav, S.P., Rothschild, G., Roumis, D.K., and Frank, L.M. (2016). Coordinated Excitation and Inhibition of Prefrontal Ensembles During Awake Hippocampal Sharp-Wave Ripple Events. Neuron 90, 113–127.

Joëls, M., and Krugers, H.J. (2007). LTP after Stress: Up or Down? Neural Plast. 2007.

Jones, M.W., and Wilson, M.A. (2005). Theta Rhythms Coordinate Hippocampal–Prefrontal Interactions in a Spatial Memory Task. PLoS Biol 3, e402.

Jung, M.W., Wiener, S.I., and McNaughton, B.L. (1994). Comparison of spatial firing characteristics of units in dorsal and ventral hippocampus of the rat. J. Neurosci. Off. J. Soc. Neurosci. 14, 7347–7356.

Kallarackal, A.J., Kvarta, M.D., Cammarata, E., Jaberi, L., Cai, X., Bailey, A.M., and Thompson, S.M. (2013). Chronic Stress Induces a Selective Decrease in AMPA Receptor-Mediated Synaptic Excitation at Hippocampal Temporoammonic-CA1 Synapses. J. Neurosci. 33, 15669–15674.

Kemere, C., Carr, M.F., Karlsson, M.P., and Frank, L.M. (2013). Rapid and continuous modulation of hippocampal network state during exploration of new places. PloS One 8, e73114.

Kim, J.J., Lee, H.J., Welday, A.C., Song, E., Cho, J., Sharp, P.E., Jung, M.W., and Blair, H.T. (2007). Stress-induced alterations in hippocampal plasticity, place cells, and spatial memory. Proc. Natl. Acad. Sci. U. S. A. 104, 18297–18302.

Kirby, E.D., Muroy, S.E., Sun, W.G., Covarrubias, D., Leong, M.J., Barchas, L.A., and Kaufer, D. (2013). Acute stress enhances adult rat hippocampal neurogenesis and activation of newborn neurons via secreted astrocytic FGF2. ELife 2.

Kitanishi, T., Ujita, S., Fallahnezhad, M., Kitanishi, N., Ikegaya, Y., and Tashiro, A. (2015). Novelty-Induced Phase-Locked Firing to Slow Gamma Oscillations in the Hippocampus: Requirement of Synaptic Plasticity. Neuron 86, 1265–1276.

Lever, C., Kaplan, R., and Burgess, N. (2014). The Function of Oscillations in the Hippocampal Formation. In Space,Time and Memory in the Hippocampal Formation, D. Derdikman, and J.J. Knierim, eds. (Vienna: Springer), pp. 303–350.

Lisman, J. (2005). The theta/gamma discrete phase code occuring during the hippocampal phase precession may be a more general brain coding scheme. Hippocampus 15, 913–922.

Luksys, G., and Sandi, C. (2011). Neural mechanisms and computations underlying stress effects on learning and memory. Curr. Opin. Neurobiol. 21, 502–508.

Mably, A.J., Gereke, B.J., Jones, D.T., and Colgin, L.L. (2017). Impairments in spatial representations and rhythmic coordination of place cells in the 3xTg mouse model of Alzheimer’s disease. Hippocampus 27, 378–392.

MacDougall, M.J., and Howland, J.G. (2013). Acute stress and hippocampal output: exploring dorsal CA1 and subicular synaptic plasticity simultaneously in anesthetized rats. Physiol. Rep. 1.

Magariños, A.M., and McEwen, B.S. (1995). Stress-induced atrophy of apical dendrites of hippocampal CA3c neurons: involvement of glucocorticoid secretion and excitatory amino acid receptors. Neuroscience 69, 89–98.

Magariños, A.M., Verdugo, J.M.G., and McEwen, B.S. (1997). Chronic stress alters synaptic terminal structure in hippocampus. Proc. Natl. Acad. Sci. 94, 14002–14008.

Makino, Y., Polygalov, D., Bolaños, F., Benucci, A., and McHugh, T.J. (2019). Physiological Signature of Memory Age in the Prefrontal-Hippocampal Circuit. Cell Rep. 29, 3835–3846.e5.

McEwen, B.S., Nasca, C., and Gray, J.D. (2016). Stress Effects on Neuronal Structure: Hippocampus, Amygdala, and Prefrontal Cortex. Neuropsychopharmacology 41, 3–23.

McHugh, T.J., Blum, K.I., Tsien, J.Z., Tonegawa, S., and Wilson, M.A. (1996). Impaired hippocampal representation of space in CA1-specific NMDAR1 knockout mice. Cell 87, 1339–1349.

Middleton, S.J., and McHugh, T.J. (2016). Silencing CA3 disrupts temporal coding in the CA1 ensemble. Nat. Neurosci.

Montgomery, S.M., and Buzsáki, G. (2007). Gamma oscillations dynamically couple hippocampal CA3 and CA1 regions during memory task performance. Proc. Natl. Acad. Sci. 104, 14495–14500.

O’>Keefe, J. (1976). Place units in the hippocampus of the freely moving rat. Exp. Neurol. 51, 78–109.

O’Keefe, J., and Burgess, N. (2005). Dual phase and rate coding in hippocampal place cells: theoretical significance and relationship to entorhinal grid cells. Hippocampus 15, 853–866.

O’Keefe, J., and Dostrovsky, J. (1971). The hippocampus as a spatial map. Preliminary evidence from unit activity in the freely-moving rat. Brain Res. 34, 171–175.

O’Keefe, J., and Nadel, L. (1978). The Hippocampus as a Cognitive Map (Oxford University Press, USA).

O’Keefe, J., and Recce, M.L. (1993). Phase relationship between hippocampal place units and the EEG theta rhythm. Hippocampus 3, 317–330.

Passecker, J., Hok, V., Della-Chiesa, A., Chah, E., and O’Mara, S.M. (2011). Dissociation of dorsal hippocampal regional activation under the influence of stress in freely behaving rats. Front. Behav. Neurosci. 5, 66.

Rahman, M.M., Callaghan, C.K., Kerskens, C.M., Chattarji, S., and O’Mara, S.M. (2016). Early hippocampal volume loss as a marker of eventual memory deficits caused by repeated stress. Sci. Rep. 6.

Resnik, E., McFarland, J.M., Sprengel, R., Sakmann, B., and Mehta, M.R. (2012). The Effects of GluA1 Deletion on the Hippocampal Population Code for Position. J. Neurosci. 32, 8952–8968.

Salehi, B., Cordero, M.I., and Sandi, C. (2010). Learning under stress: The inverted-U-shape function revisited. Learn. Mem. 17, 522–530.

Sandi, C., Davies, H.A., Cordero, M.I., Rodriguez, J.J., Popov, V.I., and Stewart, M.G. (2003). Rapid reversal of stress induced loss of synapses in CA3 of rat hippocampus following water maze training. Eur. J. Neurosci. 17, 2447–2456.

Schmitzer-Torbert, N., Jackson, J., Henze, D., Harris, K., and Redish, A.D. (2005). Quantitative measures of cluster quality for use in extracellular recordings. Neuroscience 131, 1–11.

Shin, J.D., and Jadhav, S.P. (2016). Multiple modes of hippocampal–prefrontal interactions in memory-guided behavior. Curr. Opin. Neurobiol. 40, 161–169.

Siapas, A.G., Lubenov, E.V., and Wilson, M.A. (2005). Prefrontal phase locking to hippocampal theta oscillations. Neuron 46, 141–151.

Skaggs, W.E., McNaughton, B.L., Gothard, K.M., and Markus, E.J. (1993). An Information-Theoretic Approach to Deciphering the Hippocampal Code. In In, (Morgan Kaufmann), pp. 1030–1037.

Sousa, N., Lukoyanov, N.V., Madeira, M.D., Almeida, O.F., and Paula-Barbosa, M.M. (2000). Reorganization of the morphology of hippocampal neurites and synapses after stress-induced damage correlates with behavioral improvement. Neuroscience 97, 253–266.

Suvrathan, A., Tomar, A., and Chattarji, S. (2010). Effects of chronic and acute stress on rat behaviour in the forced-swim test. Stress Amst. Neth. 13, 533–540.

Tomar, A., Polygalov, D., Chattarji, S., and McHugh, T.J. (2015). The dynamic impact of repeated stress on the hippocampal spatial map. Hippocampus 25, 38–50.

Tomar, A., Polygalov, D., Chattarji, S., and McHugh, T.J. (2021). Stress enhances hippocampal neuronal synchrony and alters ripple-spike interaction. Neurobiol. Stress 14, 100327.

Tort, A.B.L., Komorowski, R.W., Manns, J.R., Kopell, N.J., and Eichenbaum, H. (2009). Theta-gamma coupling increases during the learning of item-context associations. Proc. Natl. Acad. Sci. U. S. A. 106, 20942–20947.

Tort, A.B.L., Komorowski, R., Eichenbaum, H., and Kopell, N. (2010). Measuring Phase-Amplitude Coupling Between Neuronal Oscillations of Different Frequencies. J. Neurophysiol. 104, 1195–1210.

Vanderwolf, C.H. (1969). Hippocampal electrical activity and voluntary movement in the rat. Electroencephalogr. Clin. Neurophysiol. 26, 407–418.

Watanabe, Y., Gould, E., and McEwen, B.S. (1992). Stress induces atrophy of apical dendrites of hippocampal CA3 pyramidal neurons. Brain Res. 588, 341–345.

Yuen, E.Y., Liu, W., Karatsoreos, I.N., Feng, J., McEwen, B.S., and Yan, Z. (2009). Acute stress enhances glutamatergic transmission in prefrontal cortex and facilitates working memory. Proc. Natl. Acad. Sci. U. S. A. 106, 14075–14079.

Zaletel, I., Filipović, D., and Puškaš, N. (2016). Chronic stress, hippocampus and parvalbumin-positive interneurons: what do we know so far? Rev. Neurosci. 27, 397–409.

